# Best: A Tool for Characterizing Sequencing Errors

**DOI:** 10.1101/2022.12.22.521488

**Authors:** Daniel Liu, Anastasiya Belyaeva, Kishwar Shafin, Pi-Chuan Chang, Andrew Carroll, Daniel E. Cook

**Affiliations:** Google LLC, 1600 Amphitheatre Pkwy, Mountain View, CA

## Abstract

**Summary:** Platform-dependent sequencing errors must be understood to develop accurate sequencing technologies. We propose a new tool, `best` (Bam Error Stats Tool), for efficiently quantifying and summarizing error types in sequenced reads. `best` ingests reads aligned to a high-quality reference assembly and produces per-read metrics, summary statistics, and stratified metrics across genomic intervals. We show that `best` is 16 times faster than a prior method. In addition to being useful to support development that improves the accuracy of sequencing platforms, best can also be applied to evaluate and improve other experimental factors such as library preparation and error correction methods.

**Availability and implementation:** `best` is an open-source command-line utility available on Github (github.com/google/best) under an MIT license.

## Introduction

In recent years, a plethora of novel sequencing platforms and techniques have been developed^1–6^. Each platform, depending on its underlying technology, will produce sequence data with unique error profiles. For example, Illumina sequencing produces higher rates of mismatch errors than indel errors whereas PacBio HiFi tends to produce more errors in homopolymer regions^6,7^.

In addition to the sequencing platform, other factors contribute to the distribution and types of sequencing errors. For example sequencing errors can result from differences in library preparation^8,9^. Additionally, heuristics or machine-learning based approaches can be deployed as part of post-processing to correct sequencing errors^10,11^. To optimize sequencing accuracy, it is important that we are able to characterize the distribution and types of sequencing errors under different experimental conditions. This allows us to understand how factors such as platform, sample preparation, or post-process error correction methods impact sequencing accuracy and the types of errors impacted.

To characterize sequencing errors, a highly accurate reference assembly is required. Fortunately, the Telomere-to-telomere (T2T) consortium has developed an accurate and complete assembly of the human genome (T2T-CHM13)^12^. By aligning reads from a given sequencing platform to the T2T-CHM13 assembly, it is possible to quantify and characterize the types of sequencing errors observed for a given platform, or under different experimental conditions.

Here, we introduce a tool for characterizing sequencing errors using a reference assembly called `best`: Bam Error Stats Tool. `best` builds upon the work of a python script published in Wenger et al^6^ called bamConcordance. Although many tools exist for evaluating the quality of sequencing data^13–15^, only a few tools such as PacBio’s bamConcordance^16^ and Oxford Nanopore Technology (ONT)’s pomoxis^17^ exist for quantifying errors using a reference assembly. `best` builds upon this idea, introducing new metrics for characterizing sequencing errors and a highly efficient implementation that generalizes across different sequencing technologies.

## Implementation

`best` is a command-line tool written in Rust that quantifies sequencing errors based on alignments to a reference assembly. At its core, `best` iterates through reads aligned to a high quality reference assembly (*e.g*. T2T-CHM13), counts the number and types of errors, and aggregates these values into multiple output summary files. `best` provides a variety of outputs including quantifying errors (mismatches and indels), the yield of sequence at a given error threshold, and the distribution of indel lengths. `best` is also capable of providing error distributions stratified by read length, quality score, GC content, and other variables.

`best` also allows users to stratify errors using a custom bed file. This functionality can allow users to, for example, quantify sequence errors specific to gene elements such as coding sequences or the type of gene such as rDNA or a chromosomal region such as telomeric regions. This is also useful for evaluating regions known to affect sequencing quality, like G-quadruplex or other non-B DNA conformations^18,19^.

We chose to implement `best` in Rust for its performance and ease of parallelization. `best` uses the noodles library^20^ for reading fasta/BAM files and rayon^21^ for parallelized alignment statistics collection. This allows `best` to efficiently scale to large sequencing datasets with millions of reads.

## Application

To demonstrate the utility of `best`, we used sequence data made available from the T2T consortium that was used to generate the CHM13 assembly^12^. The T2T consortium has provided Illumina, ONT, and PacBio sequence data aligned to the CHM13 assembly that can be used to compare sequencing metrics across technologies^22^. We subsampled 10% of alignments from each BAM file and ran `best` on them. We observe that `best` scales well with more threads and that it is ~16x faster than bamConcordance when using only four threads (**Figure 1a**). We only compared `best` to bamConcordance and not pomoxis because pomoxis is implemented as a set of tools for different types of analysis rather than a single binary, and this makes it difficult to make a fair comparison on runtime.

**Figure 1.**
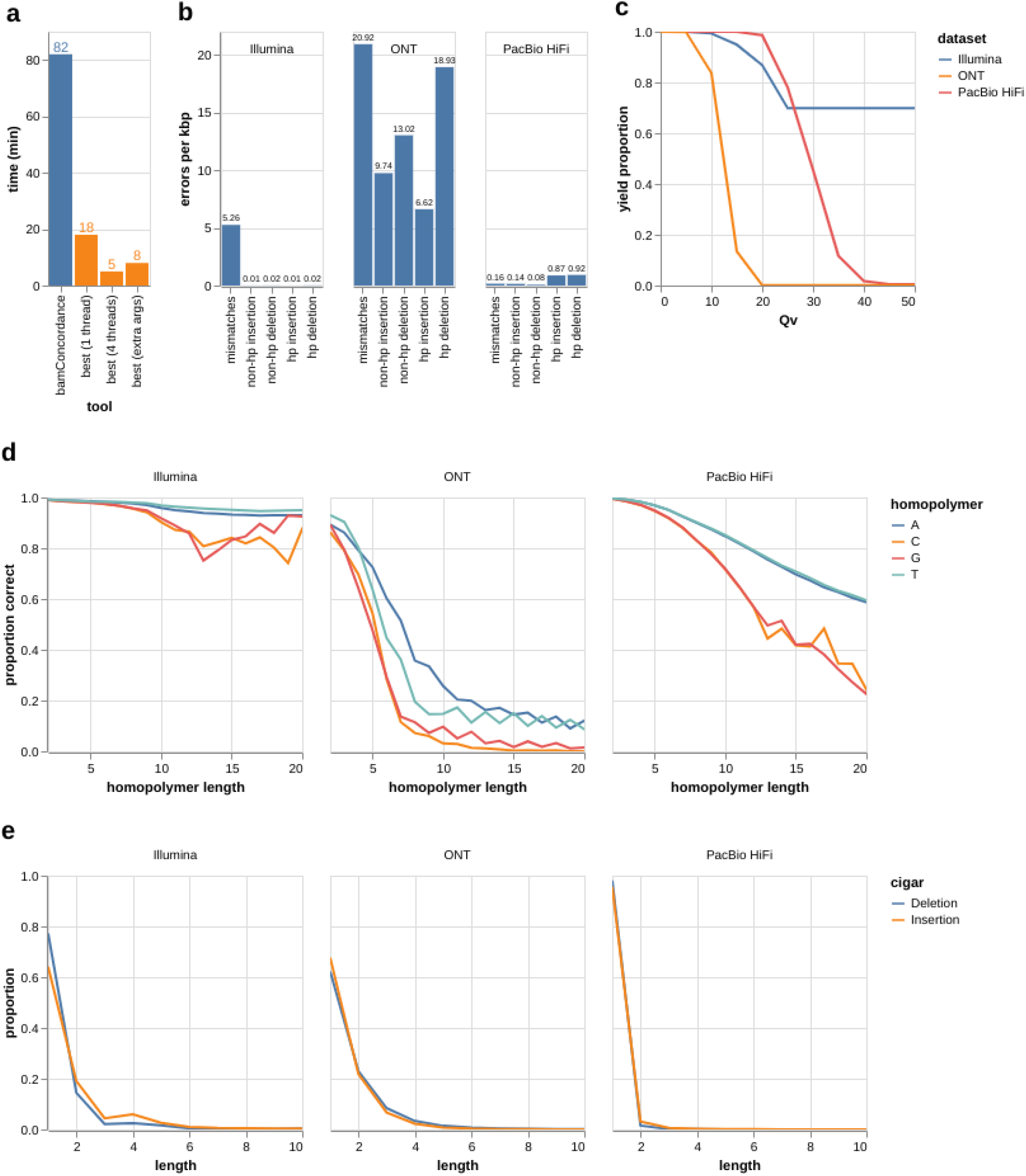
Example analysis with best. **(a)** Runtime of best with different thread counts compared to bamConcordance on 10% subsampled CHM13 PacBio HiFi data. The `best (extra args)` setup uses the following flags: `-t 4 --intervals-hp`. These flags were also used to generate plots **b-e**. **(b)** Distributions of different types of errors by platform. Insertions and deletion errors are grouped by whether they appear in the context of homopolymer (hp) or non-homopolymer (non-hp) regions. **(c)** The distribution of yield (y-axis) by empirical Qv-values (x-axis; Phred scaled). A Qv of 10 corresponds to 1 error every 10 bp, 20 = 1 error per 100 bp, etc. **(d)** The proportion of correct homopolymer regions (y-axis) by length (x-axis). **(e)** Proportion of insertions or deletions errors (y-axis) is shown by observed length of indel compared to the reference assembly (x-axis).

In **Figure 1b-1e**, we have generated plots using `best` outputs. **Figure 1b** illustrates how `best` is able to quantify sequencing errors by type and sequence context across different sequencing platforms. **Figure 1c** shows the yield curves by empirical accuracy across sequencing platforms. **Figure 1d** shows the proportion of homopolymers that are correct by homopolymer length and sequencing platform. Finally, **Figure 1e** shows the proportion of observed indel errors by length across sequencing platforms. Note that these figures are not comprehensive of the outputs made available by our tool. `best` produces additional outputs including read-level summaries, error types stratified by read length, GC content, and more that can also provide insight into the error profile of a given BAM file.

## Conclusion

`best` efficiently characterizes sequencing error profiles, providing users with insights that can be used to evaluate accuracy and help in optimizing a sequencing platform, sample preparation protocol, or error-correction process. We welcome feedback from the community, and hope that `best` will contribute to improvements in sequencing accuracy.

## References

1. Jain, M., Olsen, H. E., Paten, B. & Akeson, M. The Oxford Nanopore MinION: delivery of nanopore sequencing to the genomics community. Genome Biology vol. 17 Preprint at https://doi.org/10.1186/s13059-016-1103-0 (2016).

2. Singular Genomics. Singular Genomics https://singulargenomics.com/ (2020).

3. Website. https://www.elementbiosciences.com/r.

4. Almogy, G. et al. Cost-efficient whole genome-sequencing using novel mostly natural sequencing-by-synthesis chemistry and open fluidics platform. Preprint at https://doi.org/10.1101/2022.05.29.493900.

5. Short-read sequencing by binding. PacBio https://www.pacb.com/technology/sequencing-by-binding/ (2022).

6. Wenger, A. M. et al. Accurate circular consensus long-read sequencing improves variant detection and assembly of a human genome. Nat. Biotechnol. 37, 1155–1162 (2019).

7. Foox, J. et al. Performance assessment of DNA sequencing platforms in the ABRF Next-Generation Sequencing Study. Nat. Biotechnol. 39, 1129–1140 (2021).

8. Do, H., Wong, S. Q., Li, J. & Dobrovic, A. Reducing Sequence Artifacts in Amplicon-Based Massively Parallel Sequencing of Formalin-Fixed Paraffin-Embedded DNA by Enzymatic Depletion of Uracil-Containing Templates. Clin. Chem. 59, 1376–1383 (2013).

9. Tanaka, N. et al. Sequencing artifacts derived from a library preparation method using enzymatic fragmentation. PLoS One 15, e0227427 (2020).

10. Baid, G. et al. DeepConsensus: Gap-Aware Sequence Transformers for Sequence Correction. bioRxiv 2021.08.31.458403 (2021) doi:10.1101/2021.08.31.458403.

11. GitHub - nanoporetech/bonito: A PyTorch Basecaller for Oxford Nanopore Reads. GitHub https://github.com/nanoporetech/bonito.

12. Nurk, S. et al. The complete sequence of a human genome. Science 376, 44–53 (2022).

13. Perdomo, J. E., M. U. Ahsan, Q. Liu, L. Fang, K. Wang. A fast and flexible quality control tool for long-read sequencing data. Poster presented at: American Society of Human Genetics. in.

14. Chen, S., Zhou, Y., Chen, Y. & Gu, J. fastp: an ultra-fast all-in-one FASTQ preprocessor. Bioinformatics 34, i884–i890 (2018).

15. Babraham Bioinformatics - FastQC A Quality Control tool for High Throughput Sequence Data. http://www.bioinformatics.babraham.ac.uk/projects/fastqc/.

16. hg002-ccs/bamConcordance at master · PacificBiosciences/hg002-ccs. GitHub https://github.com/PacificBiosciences/hg002-ccs.

17. GitHub - nanoporetech/pomoxis: Analysis components from Oxford Nanopore Research. GitHub https://github.com/nanoporetech/pomoxis.

18. Weissensteiner, M. H. et al. Altered sequencing success at non-B-DNA motifs. bioRxiv 2022.06.13.495922 (2022) doi:10.1101/2022.06.13.495922.

19. Guiblet, W. M. et al. Long-read sequencing technology indicates genome-wide effects of non-B DNA on polymerization speed and error rate. Genome Res. 28, 1767–1778 (2018).

20. GitHub - zaeleus/noodles: Bioinformatics I/O libraries in Rust. GitHub https://github.com/zaeleus/noodles.

21. GitHub - rayon-rs/rayon: Rayon: A data parallelism library for Rust. GitHub https://github.com/rayon-rs/rayon.

22. GitHub - marbl/CHM13: The complete sequence of a human genome. GitHub https://github.com/marbl/CHM13.

